# Microbial Communities are Well Adapted to Disturbances in Energy Input

**DOI:** 10.1101/066050

**Authors:** Nuria Fernandez-Gonzalez, Julie A. Huber, Joseph J. Vallino

## Abstract

Although microbial systems are well-suited for studying concepts in ecological theory, little is known about how microbial communities respond to long-term periodic perturbations beyond diel oscillations. Taking advantage of an ongoing microcosm experiment, we studied how methanotrophic microbial communities adapted to disturbances in energy input over a 20 day cycle period. Sequencing of bacterial 16S rRNA genes together with quantification of microbial abundance and ecosystem function was used to explore the long-term dynamics (510 days) of methanotrophic communities under continuous versus cyclic chemical energy supply. We observed that microbial communities appear inherently well-adapted to disturbances in energy input and that changes in community structure in both treatments are more dependent on internal dynamics than on external forcing. Results also show that the rare biosphere is critical to seeding the internal community dynamics, perhaps due to cross-feeding or other strategies. We conclude that in our experimental system, endogenous feedbacks were more important than exogenous drivers in shaping the community dynamics over time, suggesting that ecosystems can maintain their function despite inherently unstable community dynamics.

**IMPORTANCE:** Within the broader ecological context, biological communities are often viewed as stable and only experience succession or replacement when subject to external perturbations, such as changes in food availability or introduction of exotic species. Our findings indicate that microbial communities can exhibit strong internal dynamics that may be more important in shaping community succession than external drivers. Dynamic ”unstable” communities may be important for ecosystem functional stability, with rare organisms playing an important role in community restructuring. Understanding the mechanisms responsible for internal community dynamics will certainly be required for understanding and manipulating microbiomes in both host-associated and natural ecosystems.

## INTRODUCTION

Microorganisms host a diverse repertoire of temporal strategies to maximize their productivity under a variety of environmental settings that undergo periodic as well as aperiodic change. Some strategies, such as circadian rhythms, require explicit molecular clocks for proper execution (1, 2), but clocks may also be present in non-photosynthetic prokaryotes (3, 4). Bacteria can also exhibit anticipatory control (5) in which they respond to external cues, such as changes in temperature and oxygen concentration (6), to predict and adapt to environmental change before it occurs. Bacteria that anticipate environmental change have an obvious fitness advantage, and anticipatory strategies may stabilize ecosystems to perturbations (7, 8). Passive temporal strategies include resource storage (9), hibernation and dormancy (10) and persister cells (11). Strategies organized over space can also increase fitness, such as diel vertical migration (12), luxury-uptake (13), chemotaxis (14) and spatially executed redox reactions via cell gliding (15) or bacterial cables (16).

Temporal and spatial strategies have largely been studied under the context of an individual’s or population’s fitness, even though such strategies can impart signatures on entire communities (17, 18), alter resource gradients that affect community function (19) and operate over a wide spectrum of scales (20). Our previous theoretical work based on non-equilibrium thermodynamics that implements the maximum entropy production (MEP) principle (21, 22) for non-steady-state systems indicates that microbial systems capable of predicting future states (either actively or passively) can acquire and dissipate more free energy than communities that lack temporal strategies (23). Likewise, communities that can coordinate function over space can increase free energy dissipation relative to non-cooperative communities (24). By accounting for spatiotemporal strategies, the analysis provides a distinction between biotic versus abiotic systems. Namely, abiotic systems maximize instantaneous free energy dissipation, while living systems utilize information stored in their genome, culled by evolution, to maximize free energy dissipation over time and space scales relevant to their predictive capabilities. Temporal strategies allow living systems under certain conditions to outperform abiotic processes, but both systems follow the same objective; they attempt to drive the system to equilibrium as fast as possible by maximizing free energy dissipation (25).

To test our MEP-based modeling approach and to explore how diverse microbial communities respond to temporally varying environments, we implemented a long-term microcosm experiment consisting of two control chemostats that received continuous input of energy in the form of methane and air, and two treatment chemostats that received periodic energy inputs by cycling the feed gas between methane plus air and just air. The modeling work based on results from this experimental system was previously described in Vallino *et al.* (2014) (26), and results indicated that temporal strategies over time scales equal to or longer than the cycle period resulted in greater energy dissipation, which were required to match experimental observations. However, this thermodynamic approach says little about the finer scale community organization that gives rise to the larger scale processes of free energy dissipation, nor the nature of the internal mechanisms that stabilize communities to external perturbations. These subjects are associated with long standing questions in ecology on the nature of community structure versus ecosystem function and stability (27).

As recently reviewed by Song *et al.* (2015) (28), as well as by Shade *et al.* (2012) (29), there are numerous definitions associated with the concept of stability that are derived from the fields of physics and engineering that are used in ecology. However, even within a single ecosystem, it is possible to have subsystems that appear unstable, while higher level components exhibit stability (30). Even the notion of stability itself is dependent on the time scale of the observation window (31). An unstable system with a millennial time scale can appear stable over an annual observation window, but unstable systems with a monthly time scale will be perceived as such over an annual window. Yet, with the development of molecular tools, several studies have now shown that microbial populations appear unstable, such as in methanogenic communities (32), phytoplankton communities (33, 34), marine sediments (35) and nitrifying bioreactors (36, 37). Many of these systems exhibited functional stability even with unstable community dynamics, while in others, ecosystem function responded to community alterations. Functional complementarity (38, 39) can explain changes in community composition for systems where exogenous drives were not, or could not be, held constant, but it is still uncertain what drives changes in community composition when exogenous drivers are constant (40, 41).

To date there has been limited research on the importance of endogenous dynamics relative to exogenous drivers on changes in community composition, but the use of microbial microcosm experiments are well suited to address these questions that can be challenging to address in field studies (42, 43). In this work, we used 16S rRNA gene sequencing together with quantification of microbial abundance and ecosystem function to explore the long-term dynamics (510 days) of a methanotrophic microbial community under both continuous and periodic energy inputs. Results suggest that microbial communities are inherently well-adapted to disturbances in energy input, with the rare biosphere critical to seeding internal community dynamics.

## RESULTS

### Ecosystem function

The experiment was divided into four phases. Initially the methane and air mixture was kept constant during Phases I and II. In Phases III and IV the chemostats were separated into control and cycled treatments, and while the control received constant energy input, cycled chemostats were subjected to cycling inputs of methane+air and air-only mixtures with a 20 day periodicity. To assess changes in ecosystem function across treatments, we characterized ecosystem processes by measuring: NH_4_^+^, NO_3_^-^+NO_2_^-^ concentrations (from here on labeled NO_3_), pH, prokaryote and eukaryote cell densities (Fig. 1); CH_4_ and O_2_ consumption and CO_2_ production rates (Fig. 2); total dissolved nitrogen (TDN), dissolved organic carbon (DOC) and nitrogen (DON), particulate organic carbon (POC) and nitrogen (PON) (Fig. S1). Gas consumption and production rates were calculated from differences in input and output gas concentrations (Fig. S2) and flow rate.

**Figure 1:**
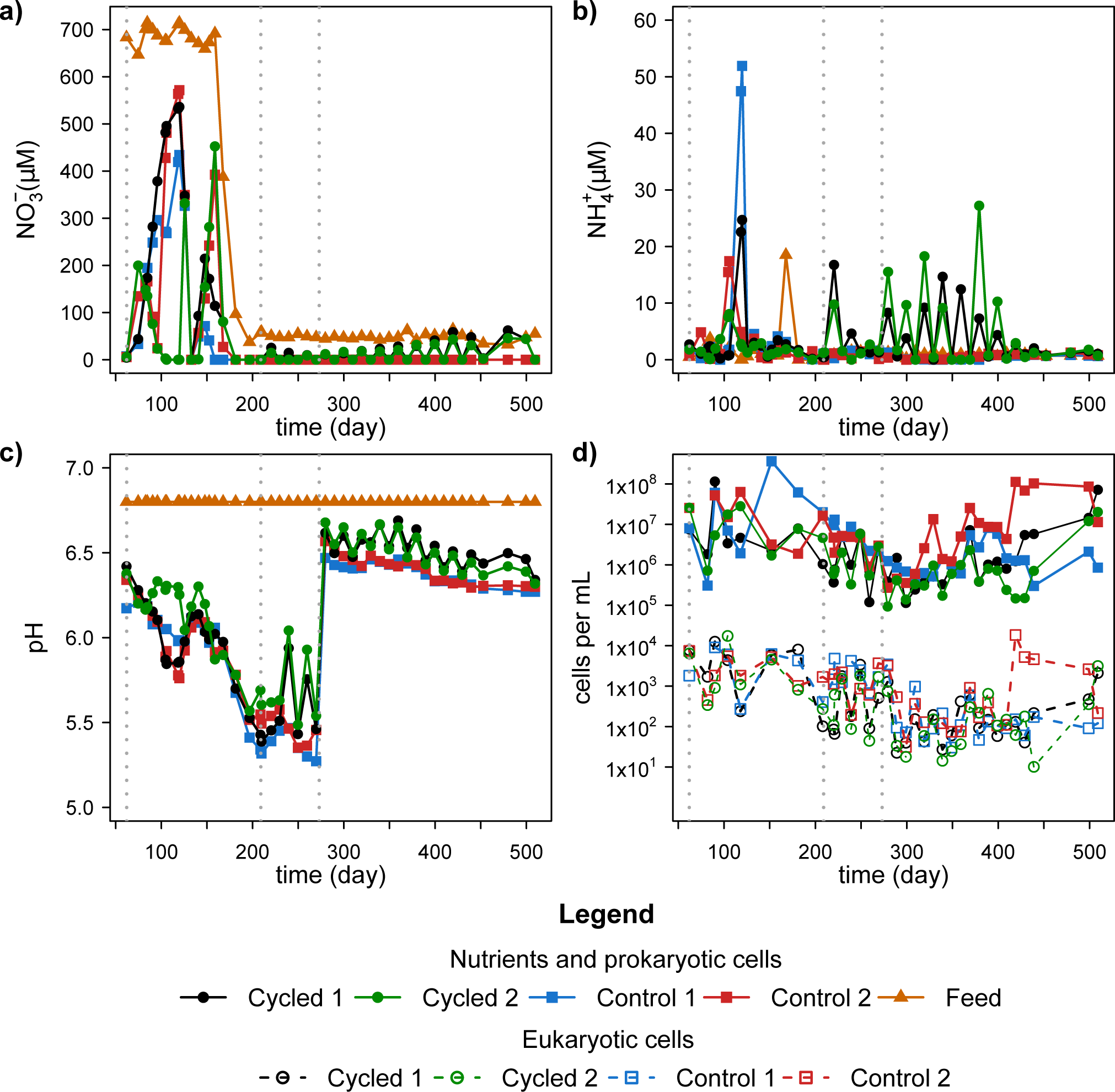
Environmental variables on chemostats and cell concentrations during Phases II, III and IV: a) nitrate+nitrite (NO_3_^-^), b) ammonium (NH_4_^+^); c) pH; d) prokaryotic and eukaryotic cell densities. Grey dotted lines indicate the start of Phases II, III and IV.

**Figure 2:**
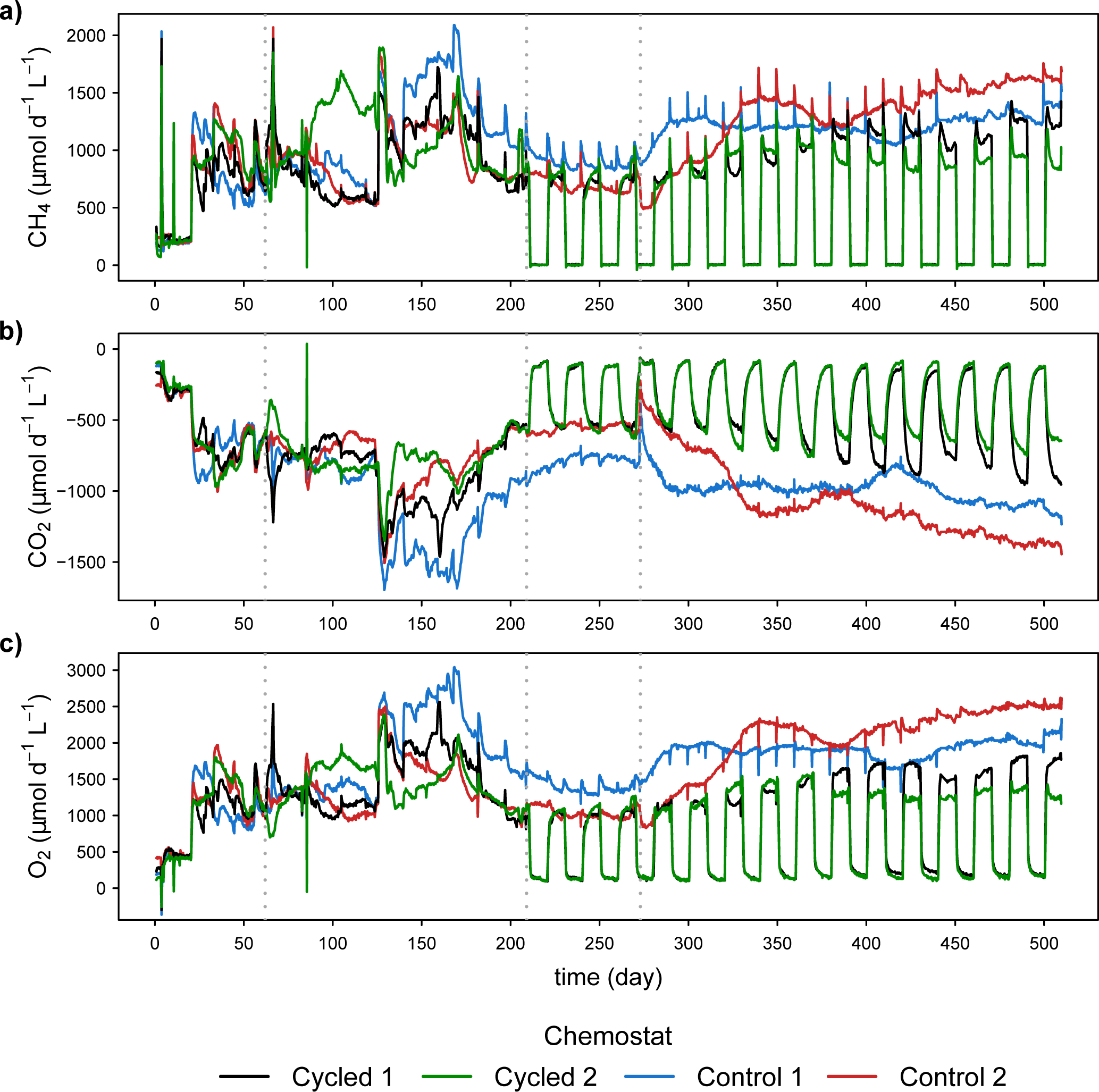
Gas production or consumption rates calculated from input and output gas concentrations and flow rate. a) Methane consumption, b) carbon dioxide production and c) oxygen consumption. Grey dotted lines indicate the start of Phases II, III and IV.

Overall, most environmental variables in the cycled chemostats exhibited the influence of periodic energy inputs during experimental Phases III and IV in the cycled chemostats, in which the measured values during CH_4_-on periods were similar to those in the control chemostats. For instance, nitrate and ammonia accumulated during the CH_4_-off periods but when CH_4_ was on, their values were drawn down almost to 0 μM, close to the control measurements (Fig. 1a, b). In the last 100 days of the experiment, periodic accumulation of ammonium was not observed even though no changes in external drivers were made in Phase IV (Fig. 1b).

Decrease in chemostats pH over Phases I, II and III (Fig. 1c) were likely caused by the increase in carbon dioxide concentration (Fig. S2) and decrease in nitrate in the feed medium (Fig. 1a). In order to minimize losses to microbial community diversity, a 10 mM, pH 6.5 phosphate buffer was added to the feed media on day 273, which defines the start of Phase IV of the experiment. All other variables, except CH_4_ feed in cycled chemostats, were maintained constant during Phase IV.

Eukaryotic and prokaryotic relative cell abundances exhibited nearly parallel behavior over the course of the entire experiment. Both cell densities fell when pH decreased in Phase III, but later recovered during the first 150 days of Phase IV (Fig. 1d). Comparing the values across treatments, no significant loss of biomass was observed in the cycled chemostats even though they only received half of the energy input compared to the controls in Phases III and IV, where microbial cell abundances were similar within and between treatments.

Gas consumption and production rates in the control chemostats showed some long-term minor fluctuations and a slow tendency to increase or decrease in Phases III and IV, but otherwise showed rather stable metabolic function (Fig. 2). In Phases III and IV, cycled chemostats gas consumption and production rates were similar, although slightly lower or higher, to those observed in the controls during CH_4_-on periods. In addition, recovery of CO_2_ gas production rates at the beginning of CH_4_-on periods were lagged compared to the CH_4_ and O_2_ rates due to carbonate chemistry dynamics, which was not accounted for in rate calculations. Changes in gas rates observed during Phases I and II were largely due to changes in operating conditions to prepare the systems for gas cycling phases.

### Effects of energy input cycling on community richness and composition

A total of 511,629 pyrosequencing sequences (7,984 – 17,288 per sample) spanning the V4V6 region of the 16S rRNA bacterial gene were clustered into 18,610 Operational Taxonomic Units (OTUs) at a 0.96 similarity cutoff (2,455 – 169 per sample). Overall, library coverage indicated that three quarters of the diversity was captured (Table S1). We did not observe any treatment effect on of bacterial richness or evenness estimations, although the values varied through time (Table S1). In particular, richness decreased and community unevenness increased in all chemostats when pH levels dropped to acidic values from late Phase II until the beginning of Phase IV (Table S1).

Morisita-Horn dissimilarity index (MH) showed that communities shifted their composition throughout the experiment with no indication of greater community similarity within treatment than between treatments (Fig. 3). A PERmutational Multivariate ANalysis Of Variance (PERMANOVA) test was used to examine if bacterial community composition within and between treatments were statistically different while accounting for the temporal trend. During Phase II when the chemostats were being mixed, communities changed in composition similarly, regardless of treatment (Table S2, Fig. 3). During both cycling Phases III and IV, bacterial community composition was more dynamic and communities changed less similarly over time (overall test, F = 1.635, P = 0.196) (Fig. 3, Table S2). The dynamic turnover of the microbial community was quite apparent when community dissimilarity between successive time points was examined in each chemostat separately (Fig. S3). Except for the start (days 62 to 104) and during the low pH Phase III (days 208 to 269), communities between two successive time points differed considerably regardless of treatment.

**Figure 3.**
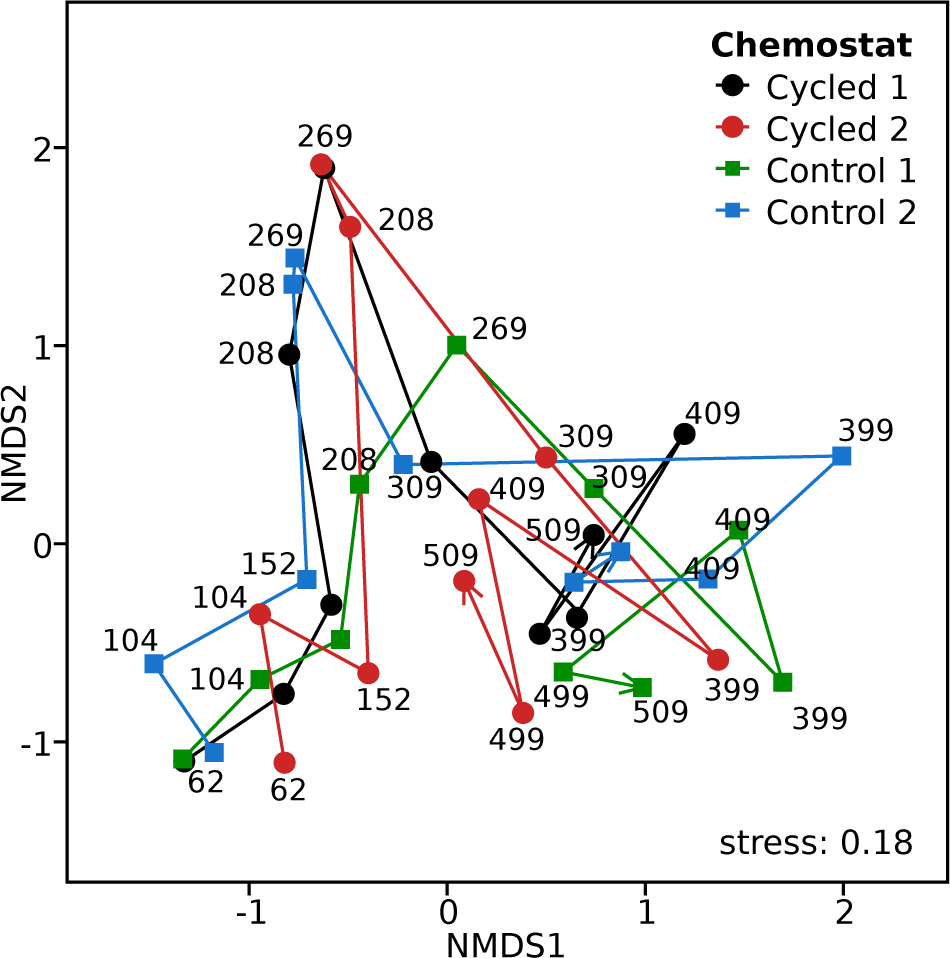
Community dissimilarities. Non-metric multidimensional scaling ordination of the Morisita-Horn dissimilarity matrix among all bacterial communities. Samples from the same chemostat are connected with arrows indicating the timeline. Labels correspond to sampling time (days).

Communities were dominated by the *Proteobacteria* phylum that averaged 70.75 and 71.47% of the community in cycled and control chemostats, respectively (Fig. S4). The most abundant class within the phylum was *Gammaproteobacteria* (44.90%, in cycled; 45.94%, in control), although classes *Alphaproteobacteria* (9.76%, 11.80%), *Betaproteobacteria* (13.90% and 12.10%), phyla *Bacteroidetes* (8.14%, 8.28%) and *Verrucomicrobia* (4.52%, 3.61%), represented a substantial percentage of communities as well.

OTUs were divided into two groups: dominant OTUs, defined as those with abundances equal to or over 1% in any of the samples analyzed in any chemostat; and rare OTUs whose abundances were always below 1%. The 150 dominant OTUs represented over 90% of the community at almost all times across all chemostats (Fig. 4, Table S3). Most of the dominant OTUs (83%) were part of the community in all 4 chemostats (Fig. S5). In addition, Linear discriminant Analysis (LDA) effect size (LEfSe) between treatments for the cycling Phases III and IV found only 18 dominant OTUs (12%), distributed differentially across control and cycled treatments (Fig. S6).

**Figure 4.**
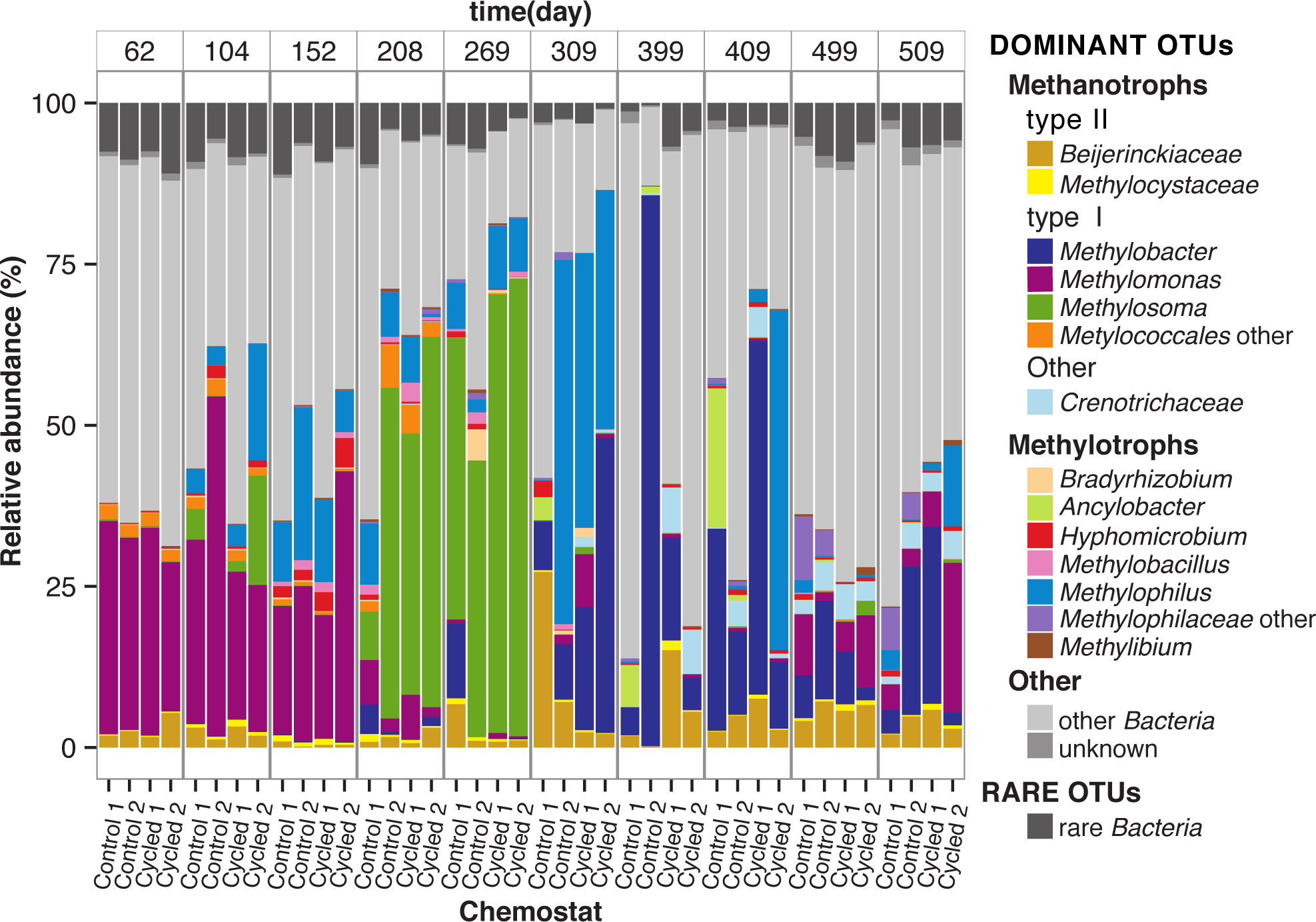
Cumulative relative abundances of dominant OTUs and rare OTUs (dark grey). For dominant OTUs, the relative abundances of one-carbon-degrading genera are indicated with colors other than grey. The “Other” category includes rest of dominant OTUs, either the remaining 66 *Bacteria* genera (Table S3) or OTUs with unknown taxonomic classification. See main text for definitions dominant and rare OTUs.

### One-carbon degrading bacteria and the rare biosphere

One-carbon (C1)-degrading bacteria, both methanotrophs and methanol degrading bacteria (hereafter referred to as methylotrophs) were a large proportion of the community at most times (Fig. 4). Those metabolic types were distributed across diverse genera, mostly within the type I methanotrophs of *Gammaproteobacteria* class (i.e. *Methylosoma*, *Methylobacter*). In addition, *Crenothrix* and putative type II methanotrophs from *Alphaprotebacteria* class (*Methylocystaceae* and *Beijerinkckiaceae* groups) were also present. Other C1 bacteria included putative methylotrophs from different families within *Betaproteobacteria* class: *Methylophilaceae* (*Methylophilus*, *Methylobacillus*) but also *Bradyrhizobiaceae* (*Bradyrhizobium*), *Hyphonomicrobiaceae* (*Hyphomicrobium* and *Ancylobacter*) and *Commamonadaceae* (*Methylibium*) (Fig. 4).

A succession of different OTUs belonging to genera in both methanotrophic and methylotrophic groups was observed over the course of the experiment (Fig. 4, Fig. 5). The C1-degrading genera successions were very similar across all chemostats during experiment Phases II and III, but some divergence was observed during pH-controlled Phase IV. Within the methanotrophs, we observed an initial dominance of *Methylomonas* sp. (mainly OTUs 13610 and 17899) during I and II, which was replaced by *Methylosoma* sp. (OTU 4859) when pH was low at the end of Phase II and Phase III (Fig. 4, Fig. 5). During Phase IV, chemostats were characterized by a more diverse and even distribution of methanotrophs, although, overall, *Methylobacter* was the most dominant genus. This phase is also characterized by a small increase in type II methanotrophs and the appearance of *Crenothrix* sp. (Fig. 4). Methylotrophs also changed with time. They were initially stimulated during early Phase II, in particular *Methylophilus* sp. (mainly OTU 15599) that was the most abundant methylotrophic genera, then replaced by *Methylobacillus* under acidic conditions (Fig. 4, Fig. 5). Later, in Phase IV, the emergence of *Ancylobacter* and unclassified *Methylophilaceae* OTUs reconfigured the assembly of methylotrophs.

**Figure 5.**
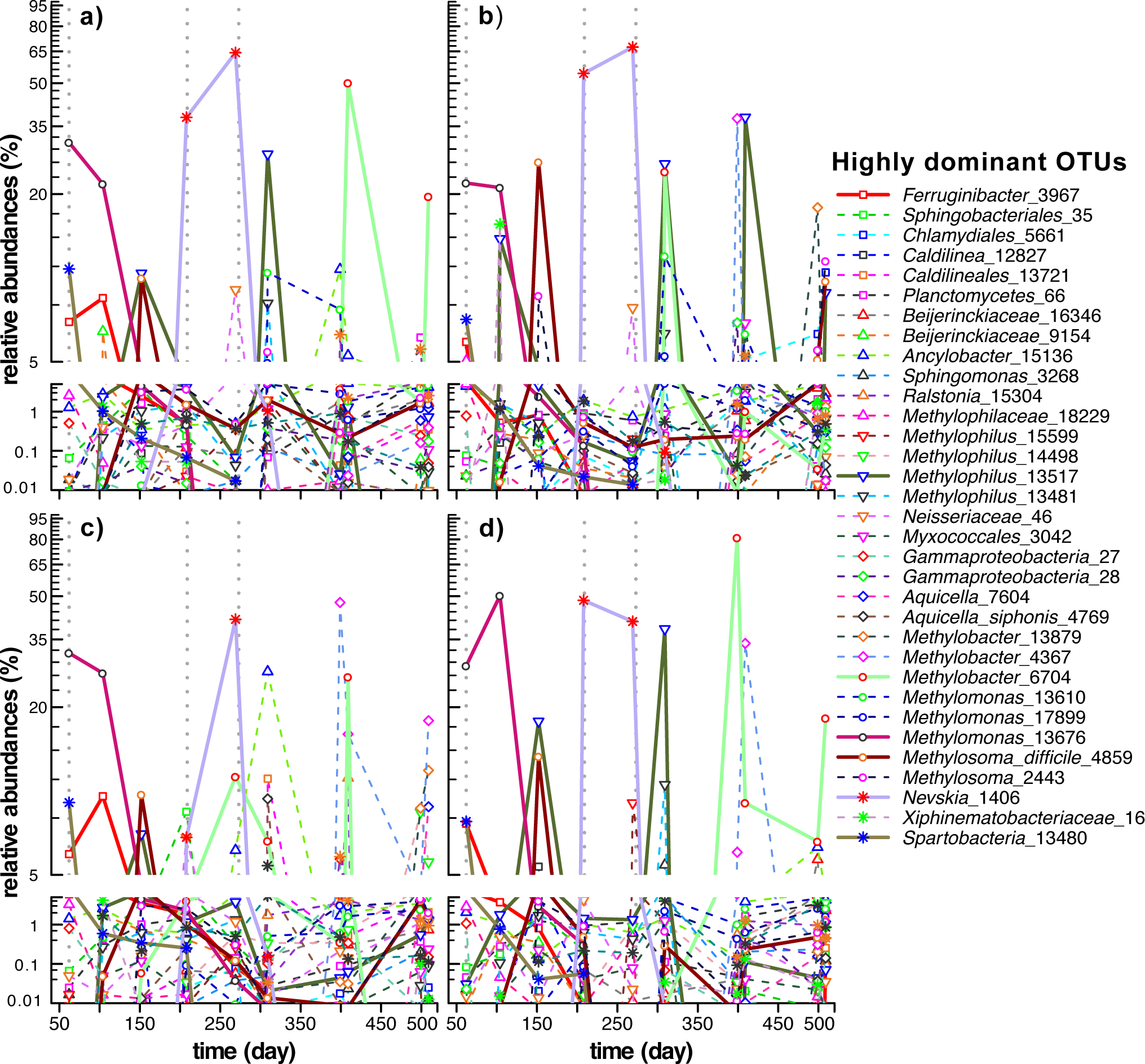
Temporal dynamics of highly dominant OTUs. Semi-logarithmic plot of temporal changes in relative abundances of dominant OTUs with relative abundances larger than 5 % in any of the samples in cycled (a, b) and control (c, d) microcosms. Solid lines identify the seven OTUs that were found at 5% or greater in all four chemostats. Names correspond to the taxonomy and number of each OTU. Note break in y-axis to highlight the importance of the rare biosphere. Grey dotted lines indicate the start of Phases II, III and IV.

In all chemostats, each one of the abundant OTUs was also member of the rare biosphere at certain times during the course of the experiment (Fig. 5). In particular, the changes in relative abundances were very large for the group of OTUs that represent more than 5% of the community in any of the samples analyzed (Fig. 5). For instance, *Methylosoma difficile*_4859 was a member of the rare biosphere on day 62 (<0.01%) in all chemostats, and at day 269, it represented over 40% of gene sequences (64%, 67%, 41%, 41% cycled-1,-2, control-1,-2 respectively). By day 399, it was back to <0.01% in all cases (Fig. 5). Other methanotrophs that were rare bacteria at the beginning of Phase II, like *Methylobacter*_13879 (<0.01% in cycled and 8x10-3 in control chemostats), was at relatively high abundances at different times (50% day 409, cycled-1; 24% day 309, cycled-2; 26% day 409, control-1 and 81% day 399, control-2). Later, this OTU behaved differently depending on the chemostat: in some cases *Methylobacter*_13879 stayed an abundant OTU for the rest of the experiment, but exhibiting large changes in relative abundance over short periods of time (9%, 7%, 18%, days 409, 499, 509 in control-2; 4%, 50%, 2%, 20% days 399, 409, 499, 509 in cycled-1). In other cases, it dropped to the rare biosphere for almost or all the rest of the experiment (8x10^-3^ %, 1%, 3x10^-2^ %, 0.4 % days 399, 409, 499, 509 in cycled-2; 0% days 499, 509 in control-1) (Fig. 5).

These types of changes were also observed for other highly abundant members of the community that are not C1-bacteria. In particular, *Aquicella siphonis*_4769 was a rare bacteria (0% in all chemostats, days 60-309) until day 399, when it represented 47%, 6% and 37% of the community in control 1, control 2 and cycled 2. Later, *A. siphonis*_4769 progressively dropped to the rare biosphere again in cycled-2 (3%, 1%, 0.01% in days 409, 499, 509), but bounced back and forth in both control chemostats (18-4% in control-1, 33-0.3% in control-2). In contrast, *A.*235 *siphonis*_4769 remained rare (0.1 – 9x10-3%) at all times in cycled-1.

## DISCUSSION

We used natural methanotrophic microbial microcosms to study how microbial communities respond to periodic inputs of energy by cycling inputs of methane and air mixtures. Overall, both the control and cycled chemostats were functionally stable. Both nitrate and ammonium concentrations increased when methane was turned off relative to the controls, but bacterial and protist cell counts were relatively stable during Phases III and IV, and there was no significant difference in cell counts between the control and cycled chemostats. Similarly, methane oxidation rate in the control chemostats during Phases III and IV was relatively constant, and while methane oxidation rate in the cycled chemostats varied as a function of the gas inputs, there were no significant changes during Phase IV. However, bacterial community composition changed in both the control and cycled chemostats during all phases of the experiment. The community dynamics in the control chemostats were particularly striking during Phase IV, given that the exogenous drivers were maintained constant during that Phase, yet the communities continued replacement of the dominant methanotroph at almost every time point. The cycled chemostats showed replacement of the dominant methanotroph at almost every sample point as well. In both control and cycled chemostats, the dominant OTU often originated from the rare biosphere (44) or was even undetected in the preceding sample.

Perhaps the most interesting result from this long term perturbation experiment was the similarity in the community dynamics between the control and cycled chemostats. We expected that the cycled chemostats would develop a dramatically different microbial community that would be better adapted to cyclic energy inputs, but the results did not support this. Instead, we observed similarity in community composition succession between the control and cycled chemostats. Considering only the most abundant OTU at each sample point, seven of them were detected across all four chemostats, and they were the most abundant OTUs in 29 out of the 40 samples examined. The speed by which the most abundant OTUs were replaced was most evident in the samples taken over a cycle period at days 399 (methane ON), 409 (methane OFF) and 499 (methane ON) and 509 (methane OFF). It may not be surprising that the most abundant OTU was replaced between methane ON (399 and 499) and methane OFF (409 and 509) in the cycled chemostats, but the switching also occurred in the controls even though methane input was constant. Furthermore, the succession in dominance was opposite in the two control chemostats at day 399 and day 409, where dominance changed from OTU-4769 to OTU-13879 in Control-1 and vice versa in Control-2. Both of these OTUs dominated in the cycled chemostats as well, with OTU-13879 dominating at end of methane OFF (day 409) and OTU-4769 dominating at the end of methane ON (day 399). From these results, it appears that endogenous feedbacks driving community dynamics were more important for shaping the community composition than exogenous drivers. Even though the cycled chemostats were significantly perturbed by periodic methane input, this exogenous forcing was of minor importance for community dynamics and composition when referenced to the control chemostats.

Our observations are similar to others. Konopka *et al.* (2007) (45) studied 16 replicate microcosms subject to discrete pulses of gelatin every day and every 7 days and observed very dynamic bacterial communities, although they observed greater variability between their replicates then we did. Similar perturbation studies (46, 47) concluded that endogenous dynamics seemed to dominate and exogenous forcing was not a strong selective pressure, which is consistent with our findings. Analysis of natural marine communities during a phytoplankton bloom also displayed rapid replacement of the dominate organisms and the importance of endogenous feedbacks in shaping communities (34). In pond microcosms, nutrient pulsing even stabilized ecosystem properties relative to non-pulsed controls via compensatory dynamics (48).

The lack of importance of exogenous drivers on community dynamics implies that the microbial communities were inherently well-adapted to periodic inputs of energy. If the microbial communities were not well-adapted to interruptions in energy availability, then we would expect that methane oxidation rates in the cycled chemostats would increase over time as the community adapted and evolved to the periodic availability of methane. This selection pressure, which was not present in the controls, would be expected to select for those organisms with enhanced resource storage capabilities that would allow growth and maintenance when methane was absent (9). Differential selection between chemostat treatments would drive changes in community composition and increases in methane oxidation rate over time. But neither of these outcomes were observed, which leads us to conclude that the communities were already well-adapted to interruptions in energy input; there was little differential selection between chemostats because effective temporal strategies were already present in both treatments. Our previous modeling work (26) also supports the conclusion that the communities were well-adapted to periodic inputs of energy, because the thermodynamically-based optimal allocation model was only able to accurately simulate the observed methane oxidation rates in the control and cycled chemostats when the optimization interval (i.e., time scale of the implied temporal strategies) was set to be equal to or greater than the 20 day methane cycle period. When shorter optimization intervals were used, the model was unable to fit the observations (see Table 18.2 in (26)). Observations near the end of the experiment (day 1242, data not shown) indicate that the temporal strategies were not clock based, because oscillations were not observed in gas dynamics in the cycled chemostats when the methane cycling was stopped. Such residual oscillations are observed in clock-based circadian systems when exogenous cycling is terminated (49). The lack of residual oscillations when methane was left ON indicates the communities probably implemented passive temporal strategies, such as resource storage, which have been identified in methanotrophs that are known to store polyhydroxybutyrate under cyclic inputs (50) as well as store fatty acids (51). If the exogenous drivers were not responsible for the observed community dynamics in both the control and cycled chemostats, what might explain the endogenous dynamics?

Changes in community composition are often associated with changes in exogenous drivers as explained by functional complementarity (38) or compensatory dynamics in response to press or pulse perturbations (39). These theories have been put forth to explain biodiversity maintenance and why competitive exclusion (52) does not lead to the “Paradox of the Plankton” (53). In complementarity, each species has evolved to grow maximally under a narrow set of environmental conditions, such as pH, temperature, light level, etc. As external drivers change the environment, such as lowering pH, succession in community composition follows, where those organisms that optimally match the new conditions are selecting for, provided sufficient biodiversity exists within the system or can be readily imported by transport processes. The rare biosphere can serve as reservoir for organisms whose traits are currently suboptimal under the prevailing conditions (41). Functional complementary and similar theories may indeed explain community succession we observed during Phases I-III in both control and cycled chemostats due to changes in pH and nutrient concentrations, where the ecosystem function of methane oxidation was maintained relatively stable by a succession of optimally adapted OTUs. Ideas derived from complementarity have also been exploited by trait-based modeling approaches (54). However, complementarity or compensatory dynamics do not explain the observed succession in OTUs during Phase IV of our experiment where external drivers were constant.

Theories such as niche complementarity, niche construction, resource partitioning, cross-feeding and others have been proposed to explain endogenous dynamics that occur in the absence of exogenous forcing (40). A basic premise in these theories is that organisms modify their environment thereby creating new niches that can be exploited by others, which can lead to natural and perpetuating succession of organisms that can occur rapidly in microbial systems (55–57). One type of niche creation is known as cross-feeding, in which the waste products of one organism’s metabolism become the food for another. In the original study by Rosenzweig *et al.* 1999 (58), cross-feeding developed from a clonal population of *Escherichia coli* that oxidized glucose completely to CO_2_, but after more than 700 generations stable polymorphisms evolved that produced and consumed acetate and glycerol intermediates (also see (59)). Hence, the clonal population naturally evolved a type of distributed metabolism (60). Syntrophy and resource partitioning are also examples of cross-feeding that develop between species, phyla and domains (61, 62), where differential production of shared intermediates over time can give rise to asymmetric population dynamics that can stabilize ecosystem function (63, 64). One mechanism that may drive evolution of cross-cross feeding is the interplay between growth rate and growth efficiency.

Metabolic analysis in substrate limited systems has shown that metabolic pathway truncation, such as partial oxidation of glucose to acetate instead of CO_2_, can result in faster energy extraction per unit time, which can support faster growth rates but leads to the excretion of by-products, such as acetate (65, 66). Accumulation of intermediates can then foster adaptive gene loss that reinforces cross-feeding (67, 68). Leaking substrates is in contrast to conventional wisdom that considers maximizing substrate use efficiency a virtue. However, a recent modeling study by González-Cabaleiro et al. 2015 (69) examined these tradeoffs and showed that maximizing energy harvest rate from substrates with its attendant by-product production accurately predicts the distribution of metabolic labor observed in multi-step anaerobic fermentation of glucose to methane and CO_2_, two-step aerobic autotrophic nitrification, and single-step anaerobic denitrification. Furthermore, excreted substrates in communities can drive the production of new metabolites that are otherwise not produced when organisms are grown in isolation via emergent biosynthetic capacity (70). Consequently, we speculate that a potential driver of the rapid succession of dominant OTUs observed in both the control and cycled chemostats may be the result of continuous niche reconstruction via the extracellular accumulation of metabolic intermediates. As intermediate metabolites accumulate beyond certain thresholds, select members from the rare biosphere may be freed from dormancy by competitive advantages that allow them to achieve dominance, but only temporarily. New dominant OTUs might excrete new intermediate metabolites that then eventually select for new replacements. With sufficient biodiversity, intermediate metabolites come and go, but none accumulate significantly, so ecosystem function, such as methane oxidation rate or primary production, proceed at maximum despite the continuous species turnover. Of course, in some situations, violent perturbations can lead to excessive accumulation of metabolites and cause system collapse (71).

Dynamic cross-feeding is not the only process shaping communities. Depending on the characteristic time scales of endogenous and exogenous forcings, generalist and specialist also arise (72) and various types of chemical warfare are likely at play (73). Cooperation via cross-feeding (74, 75), quorum sensing (76), stigmergy (77), horizontal gene transfer (78) and other types of intercellular communication (79, 80) also contribute to endogenous feedbacks that likely support the continuous succession of dominant OTUs we observed. Indeed, the continual turnover of the community may be a significant mechanism in producing and maintaining the rank abundance distribution of the rare biosphere (44). Furthermore, the endogenous dynamics exhibited by microbial systems brings into question the usefulness of stability criteria often used to assess and cull food web models. If microbial community dynamics are fundamentally unstable (81), then predicting ecosystem function based on maximizing dissipation of free energy may be more tractable approach for understanding how communities will change to exogenous forcings (23).

Our results, as well as results from a previous modeling study (26), indicate that the microbial communities in our methanotrophic microcosms are inherently well-adapted to periodic inputs of energy, likely due to implementation of temporal strategies, such as resource storage. The 16S rRNA gene sequences were sampled deeply at 10 time points during the 514 day experiment and show that the dominant OTU at any time point often originated from the rare biosphere, but was subsequently replaced by a new competitor also derived from the rare biosphere at the next sample point. Even though the control and cycled chemostats experienced significantly different exogenous forcing and overall community composition changed as the experiment proceeded, the succession in dominant OTUs in both treatments were more similar than different. These results indicate that endogenous feedbacks were more important than exogenous drivers in shaping the community dynamics over time. Based on literature support, we speculate that dynamic cross-feeding may be the mechanism given rise to the unstable community dynamics. Furthermore, our results, as well as others, bring into question the usefulness of the stability concept for understanding food web architectures in microbial systems. Because ecosystem function, namely methane oxidation rate, was insensitive to the community dynamics, our results support the conjectures that systems organize to maximize free energy dissipation that many different food web configurations can support as evident by the observed community succession.

## MATERIALS AND METHODS

### Experiment set up and sampling

The experimental set up consisted of four 18 L Bellco Glass bioreactors housed in a dark Conviron environmental chamber controlled at 20°C. The microcosms were previously inoculated with 1 L of unfiltered water collected from a cedar bog in Falmouth (Massachusetts, USA) and sparged with a gas mix containing 4.9% methane in air at a gas flow rate of 20 mL min^-1^ (for details see supplementary materials and methods). The experiment consisted of four phases (Fig. 6). Phase I (days 0-62): microcosms initially operated in batch mode, but all reactors were interconnected in a closed loop at a flow rate of 10 mL min^-1^ to insure uniform community composition between MCs. Phase II (days 63-209): MCs operated independently in chemostat mode with a defined mineral salt medium (70 μM K_2_HPO_4_, 700 μM KNO_3_, 100 μM MgSO_4_, 100 μM CaCl_2_, 100 μM NaCl) plus trace elements (final concentrations: 18.50 μM FeCl_3_ 6H_2_O, 0.49 μM H_3_BO_3_, 0.13 μM CoCl_2_ 6H_2_O, 0.10 μM CuSO_4_ 5H_2_O, 0.35 μM ZnSO_4_ 7H_2_O, 0.16 μM MnSO_4_ H_2_O, 0.12 μM Na_2_MoO_4_ 2H_2_O, 0.08 μM NiCl_2_ 6H_2_O, 0.1 mM HCl) at dilution rate of 0.1 d^-1^ (1.25 mL min^-1^). Nitrate concentration was adjusted decrementally from 700 μM to 50 μM to ensure N-limited rather than CH_4_-limited growth. Phase III (days 210-273), duplicate chemostats were divided into control and cycled treatments. Cycled chemostats were subject to periodic energy input cycles by switching gas composition from methane (4.9%) plus air mixture to solely air on a 20 day period (i.e., 10 days CH_4_-on, 10 days CH_4_-off). The two control chemostats were maintained under continuous input of 4.9% methane in air (Fig. 6). Phase IV (days 274-510): gas cycling continued but passive pH control was initiated by adding 10 mM potassium phosphate buffer in the feed medium. Liquid samples were withdrawn periodically for analysis of nitrate+nitrite (NO_3_^-^), ammonia (NH4^+^), particulate organic carbon (POC) and nitrogen (PON), dissolved organic carbon (DOC) and nitrogen (DON) and microbial cell abundances (both eukaryotic and prokaryotic organisms). CH_4_, O_2_, CO_2_ gas concentrations in the feed and headspace were automatically measured and recorded every 5 hours, and pH was recorded every hour. For details on the analysis see supplementary materials. Biomass samples for 16S rRNA gene sequencing were taken on 10 different days, where all samples except for days 399 and 499 correspond to periods when CH_4_ was on in the cycled chemostats (Fig. 6). Up to 600 mL were filtered through 0.22 μm Sterivex-GP membranes and immediately frozen at -80 °C until DNA extractions.

**Figure 6.**
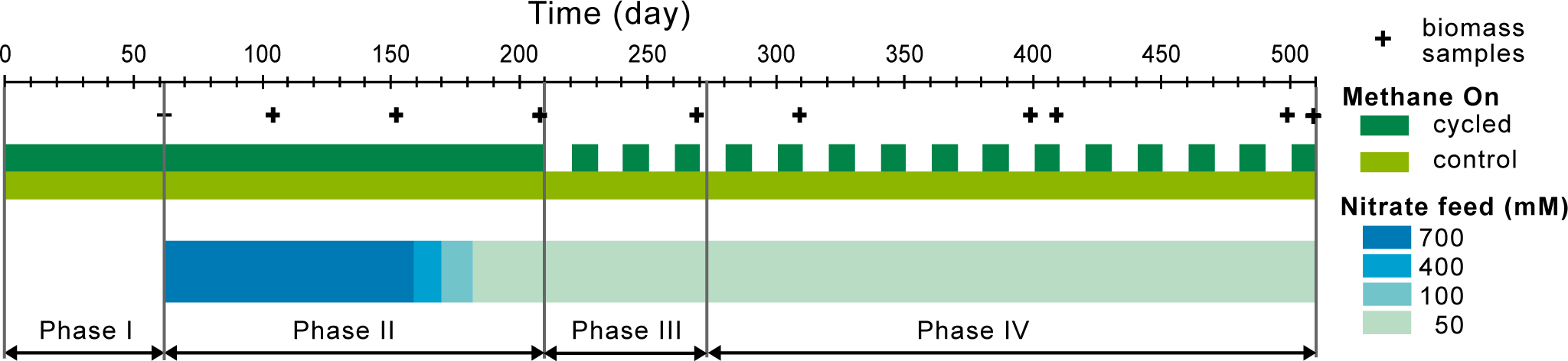
Experiment timeline showing the different phases of the study, the methane presence or absence in the gas feed for each treatment, the changes in nitrate concentrations in the liquid feed and the sampling dates for microbial community characterization. Phase I: batch-mode, Phase II: start-up, Phase III: cycling, Phase IV: pH-controlled cycling.

### DNA extraction, pyrosequencing and sequence analysis

Total genomic DNA was extracted from Sterivex filters that were thawed and cut into small pieces prior to extraction of nucleic acids using RNA PowerSoil^®^ Total RNA Isolation kit combined with the DNA Elution Accessory kit (MoBio, Carlsbad, CA, USA) following the manufacturer’s protocol. DNA concentrations were determined with Quant-iT PicoGreen dsDNA Assay Kit (Life technologies, Grand Island, NY, USA). Amplicon libraries for the V4V6 region of 16S rRNA bacterial genes were prepared using fused primers and sequenced in a Roche Titanium as previously described (82). Sequencing reads were quality filtered to remove any reads containing ambiguous bases, with average quality scores below Q30 or lacking exact primer matches. Quality-filtered sequences were analyzed for chimera removal with Uchime (83), combining both the novo and reference database (ChimeraSlayer GOLD) modes and then, clustered at 0.96 similarity with Uclust (84) to define OTUs. Taxonomy was assigned by global alignment for sequence taxonomy (GAST; Huse et al., 2008) with the SILVA database (86). Quality-filtered sequences are publicly available through the VAMPS database (https://vamps.mbl.edu) under the project JAH_ENT_Bv6v4. Raw reads for V6 are available in the NCBI Short Read Archive under Accession Number PRJNA322031.

Statistical analysis of OTUs abundances was performed with QIIME 1.8 (87), PRIMER 6 and PERMANOVA+ (Primer-E Ltd., Plymouth, United Kingdom) (88) and R (89). To compare bacterial communities and estimate community turnover, a distance matrix was calculated using the Morisita-Horn dissimilarity index (MH) (90) of log-transformed rarefied data. Non-Metric Multidimensional Scaling (NMDS) analysis was applied to explore distances among communities. Differences between treatments (control and cycled) and time (sampling day) were tested with PERMANOVA tests (91) with 1000 replications including pair-wise comparisons between individual samples.

## ACKNOWLEDGMENTS

We thank Chris Algar, Holly Cantin, Richard Fox, Hilary Morrison, Emily Reddington, and Stephanie Strebel for experimental, intellectual, and facility support. We are grateful for support from the National Science Foundation (Grants EF-0928742 to JJV and JAH and OCE-1238212 for JJV).

## SUPPLEMENTARY MATERIAL

**Supporting information**. Supplementary materials and methods.

**Figure S1.** Environmental variables on microcosms and media feed for Phases II, III and IV; a) dissolved organic carbon (DOC), b) dissolved organic nitrogen (DON) and c) total dissolved nitrogen (TDN). Grey dotted lines indicate the start of Phases II, III and IV.

**Figure S2.** Gas concentrations: a) methane, b) carbon dioxide and c) oxygen. Grey dotted lines indicate the start of Phases II to IV.

**Figure S3:** Community turnover. Morisita-Horn dissimilarities between communities from consecutive days within each microcosm. Grey dotted lines indicate the start of Phases II to IV.

**Figure S4:** Community composition at high taxonomic level. Relative abundances of phyla for each microcosm through time. *Proteobacteria* phylum has been divided into its classes. Phyla *Proteobacteria*, *Bacteroidetes*, *Verrucomicrobia*, *Acidobacteria*, *Chloroflexi*, *Planctomycetes*, *Chlorobi*, *Cyanobacteria*, *Actinobacteria*, OP10, *Nitrospira*, *Gemmatimodadetes*, TM7, *Firmicutes*, BRC1, *Spirochaetes*, *Lentisphaerae*, *Fibrobacteres* and OD1 are commonly found on freshwater environments.

**Figure S5.** Venn diagram of shared dominant OTUs between all microcosms during Phases all experimental phases.

**Figure S6**: Relative abundances of statistically significant differentiating OTUs for each treatment during phases III and IV based on LEfSe results. OUT label color indicates the treatment for which it is significant; cycled: green, control: blue. Data is averaged by day.

**Table S1:** Number of sequences, library coverage and alpha diversity indices.

**Table S2:** PERMANOVA pairwise tests results.

**Table S3:** Relative abundances of lower taxonomic groups that contains abundant OTUs.

